# The Genome of the Mustard Hill Coral, *Porites astreoides*

**DOI:** 10.1101/2022.07.01.498470

**Authors:** Kevin H. Wong, Hollie M. Putnam

## Abstract

Coral reefs are threatened both locally and globally by anthropogenic impacts, which to date have contributed to substantial declines in coral cover worldwide. However, some corals are more resilient to these environmental changes and therefore have increased relative abundance on local scales and may represent prominent members shaping future reef communities. Here, we provide the first draft reference genome for one such reef-building coral, the mustard hill coral, *Porites astreoides*. This reference genome was generated from a sample collected in Bermuda, with DNA sequenced via Pacific Biosciences HiFi long-read technology to provide an initial draft reference genome assembly. Assembly of the PacBio reads with FALCON UnZip resulted in a 678 Mbp assembly with 3,051 contigs with an N50 of 412,256. The genome BUSCO completeness analysis resulted in 90.9% of the metazoan gene set. An *ab initio* transcriptome was also produced with 64,636 gene models with a transcriptome BUSCO completeness analysis of 77.5% when compared to the metazoan gene set. The function annotation was obtained through a hierarchical approach of SwissProt, TrEMBL, and NCBI nr database of which 86.6% of proteins were annotated. Through our *ab initio* gene prediction for structural annotation and generation of a functional annotation for the *P. astreoides* draft genome assembly, we provide valuable resources for improving biological knowledge, which can facilitate comparative genomic analyses for corals, and enhance our capacity to test for the molecular underpinnings of adaptation and acclimatization to support evidence-based restoration and human assisted evolution of corals.

**Classifications:** Genetics and Genomics; Animal Genetics; Marine Biology

## Data Description

### Context

Coral reef ecosystems provide disproportionately large economic, scientific, and cultural value for their global footprint. These ecosystem engineers build coral skeletons that are home to a variety of marine life. This engineering capacity is due to the nutritional symbiosis between the coral host and their dinoflagellate algal symbionts, Symbiodinaceae (LaJeunesse et al. 2018), which provides excess carbon to fuel coral metabolic processes (Muscatine and Cernichiari 1969) and stimulate growth of the 3D skeletal structure. Under thermal stress as little as 1°C above local summer maxima there is a breakdown in the symbiosis of corals and Symbiodinaceae (Coles and Brown 2003), which can result in mass mortality that can reshape reef communities (Hughes et al. 2021).

In the face of rapid climate change, reef-building corals have been under unprecedented stress resulting in global population declines of these fundamental species (Hughes et al. 2018). While some species are sensitive to climate change (i.e., ecological losers), others are more resistant or resilient under a rapidly changing environment (i.e., ecological “winners”) (Loya et al. 2001). These taxa are often able to grow and reproduce in a “weedy” fashion, and thus increase in relative abundance on the reef (Darling et al. 2012). One such weedy coral species is the mustard hill coral, *Porites astreoides* (Lamarak, 1826), which is a ubiquitous shallow Western Atlantic coral that is present across large environmental clines from mesophotic depths of ∼45m (Goodbody-Gringley et al. 2021) to shallow mangrove environments (Lord et al. 2021) and a latitudinal range from Brazil (Pereira et al. 2017) to Bermuda (Dodge, Logan, and Antonius 1982). In contrast to other Caribbean corals, *P. astreoides* has increased in abundance in recent years, with high juvenile (Vermeij et al. 2011) and adult (Green, Edmunds, and Carpenter 2008; Toth et al. 2019; Eagleson et al. 2021) abundances, but smaller colony sizes (Edmunds, Didden, and Frank 2021). *P. astreoides* is considered a “weedy species’’ as it is a hermaphroditic, brooding coral species that has a prolonged planulation period (Chornesky and Peters 1987; McGuire 1998) with pelagic larvae that exhibit high phenotypic plasticity (de Putron and Smith 2011; de Putron et al. 2017; Goodbody-Gringley et al. 2018; Zhang et al. 2019; Wong et al. 2021) and high recruitment rates (Goodbody-Gringley et al. 2018, 2021). Additionally, larvae and juvenile life stages have been well studied under ocean acidification (Albright, Mason, and Langdon 2008; de Putron et al. 2011) and temperature stress condition (Edmunds, Gates, and Gleason 2001; Edmunds et al. 2005; Ross et al. 2013; Olsen et al. 2013; Zhang et al. 2019) due to the ease of brooded larval collection. Adult colonies also display phenotypic plasticity in response to different reef environments (Elizalde-Rendón et al. 2010; Kenkel et al. 2013; Kenkel, Meyer, and Matz 2013; Haslun et al. 2018; Walker et al. 2019; Lenz et al. 2021; Lord et al. 2021; Wong et al. 2021) and recovery from thermal bleaching events (Kenkel et al. 2013; Kenkel, Meyer, and Matz 2013; Grottoli et al. 2014; Schoepf et al. 2015; Kenkel, Setta, and Matz 2015; Levas et al. 2018; Wong et al. 2021; Bove et al. 2019; Chapron et al. 2022).

As ‘omics approaches have emerged, there has been an increase in the study of *P. astreoides* for genetic connectivity and population structure (Serrano et al. 2016; Gallery et al. 2021; Riquet et al. 2022), microbiome community (Wegley et al. 2007; Thurber et al. 2012; Sharp, Distel, and Paul 2012; Meyer, Paul, and Teplitski 2014; Glasl, Herndl, and Frade 2016; Fuess et al. 2017; Meiling et al. 2021; Dunphy, Vollmer, and Gouhier 2021), Symbiodinaceae community (Reich, Robertson, and Goodbody-Gringley 2017; Serrano et al. 2016; Salas et al. 2017), gene expression (Kenkel, Meyer, and Matz 2013; Fuess et al. 2017; Walker et al. 2019; Mansour et al. 2016), and epigenetics (Dimond and Roberts 2020). Although three *de novo* transcriptome assemblies are available for this species (Kenkel, Meyer, and Matz 2013; Mansour et al. 2016; Walker et al. 2019) they have relatively low coverage of the anticipated gene repertoire (e.g., BUSCO scores are only 18.1-26.5% complete with respect to the single copy metazoan reference gene set). Therefore, the field is currently limited by the lack of an available reference genome and improved transcriptome. These resources would greatly enhance studies that are reliant on a reference genome, for example Whole Genome Bisulfite Sequencing and Genome-Wide Association Studies. Our study is the first to generate a publicly available, assembled, and structurally and functionally annotated reference genome of *P. astreoides*, in addition to an improved reference transcriptome.

## Methods

### Coral collection, treatment, and sampling

One adult *P. astreoides* colony (Figure 1A) was collected on June 12, 2017 from Bailey’s Bay Reef Flats (32°22′27″N, 64°44′37″W) in Bermuda, transported to the Bermuda Institute of Ocean Sciences. The colony was fragmented into genetic replicates using a drill and 3.5 cm diameter size hole saw to generate circular cores of tissue and skeleton (Figure 1B). Replicate fragments were affixed to plugs using underwater epoxy (HoldFast Epoxy Stick, Instant Ocean), which covered all of the skeletal surface. The replicates fragments were held in indoor tanks with flowing seawater with LED lights (Arctic-T247 Aquarium LED, Ocean Revive) under ambient conditions for 14 days (28°C with 12:12hr light cycle at ∼115 µmol photons) and then exposed to either ambient or heated conditions (31°C with 12:12hr light cycle at ∼115 µmol photons). These conditions were applied for 59 days in order to reduce the concentration of endosymbiotic dinoflagellates (Symbiodiniaceae) and thereby to enrich for host DNA for downstream DNA extraction and sequencing. Coral fragments were immediately snap frozen in liquid nitrogen and stored at -80°C on August 28, 2017. An additional four fragments were sampled for RNA sequencing. Of these fragments, two were fragments under ambient conditions, one fragment that experienced the 59 day thermal stress with an additional hyposalinity stress (approx. 18 psu) 30 minutes before snap freezing, and one fragment under ambient conditions from a different *P. astreoides* colony from the same reef site.

**Figure 1.**
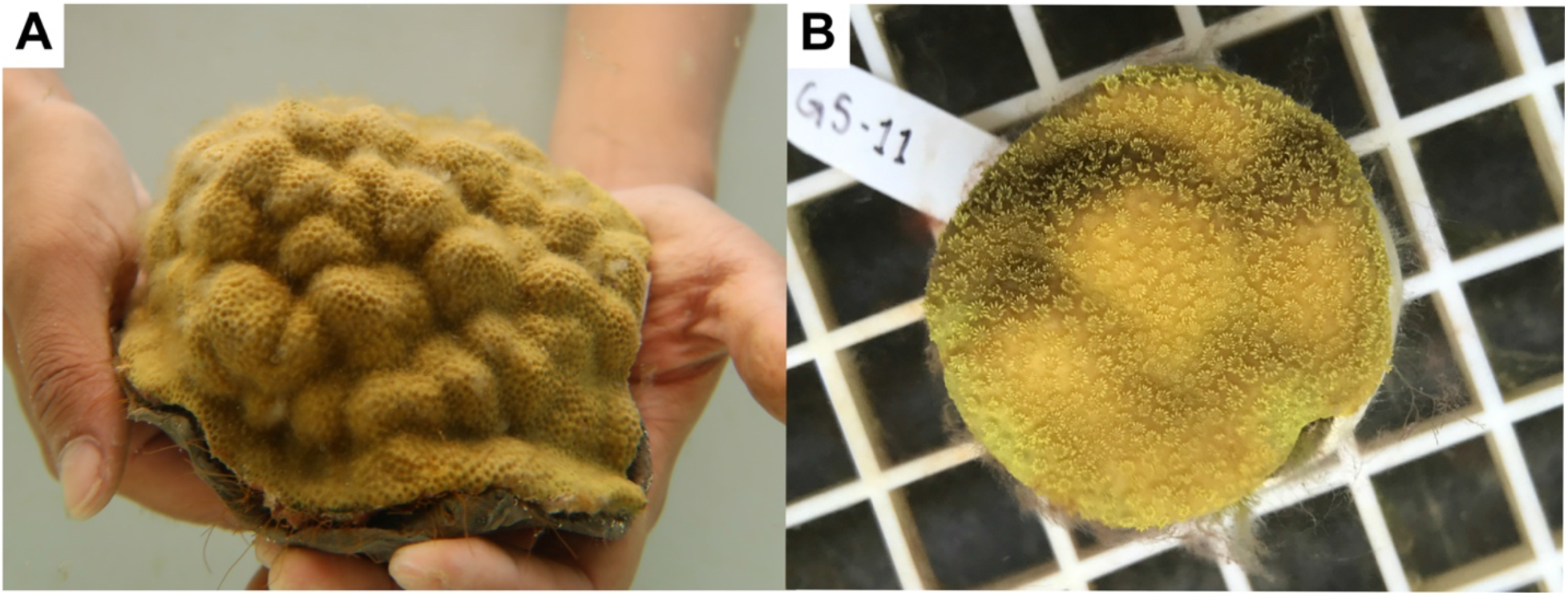
A) Colony of *P. astreoides* and B) replicate fragment cores used for exposures and extractions.

### Extraction of Genomic DNA

The frozen coral samples were homogenized with a mortar and pestle and liquid nitrogen. Coral high molecular weight genomic DNA (gDNA) was extracted using the QIAGEN Genomic-tip 100/G (Cat #10223), the QIAGEN Genomic DNA Buffer Set (Cat #19060), QIAGEN RNase A (100mg/mL concentration; Cat #19101), QIAGEN Proteinase K (Cat #19131), and DNA lo-bind tubes (Eppendorf Cat #022431021) according to the manufacturer’s instructions for preparation of tissue samples in the QIAGEN Genomic DNA Handbook (version 06/2015) using 5 individual sample preps of the same homogenized material to maximize yield. gDNA concentration was quantified using the Qubit DNA Broad Range kit (Invitrogen, Cat #Q32850) and DNA integrity was measured using a Genomic D5000 Screentape on an Agilent TapeStation 4200 system (Figure 2).

**Figure 2.**
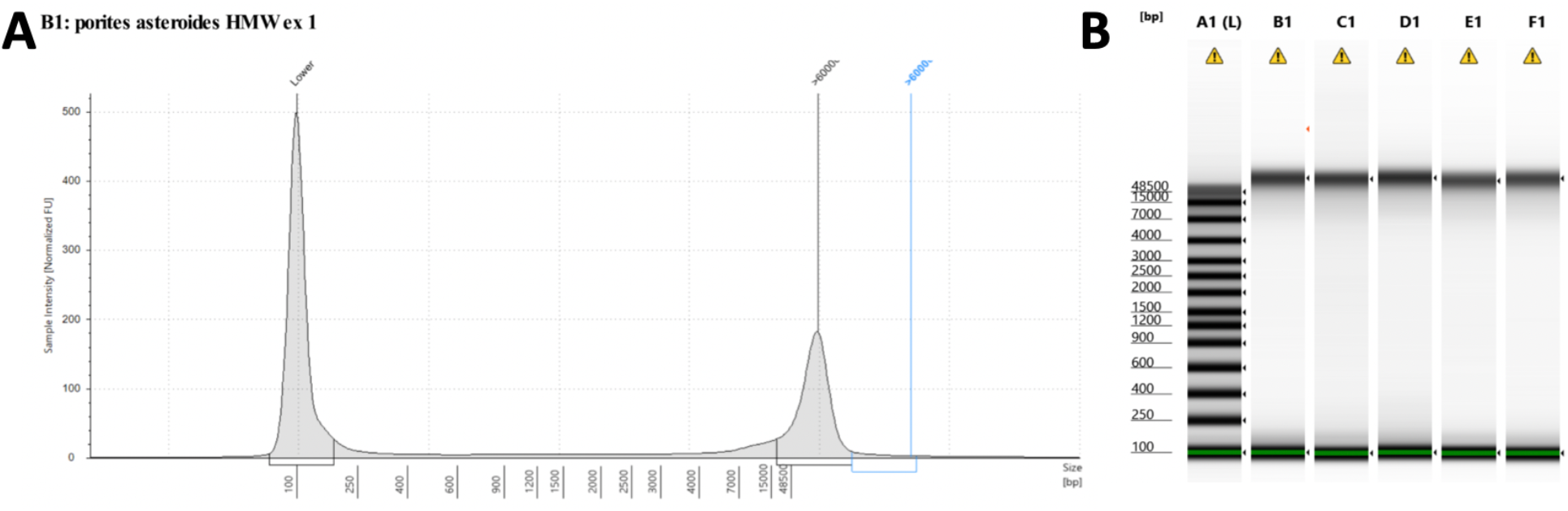
Quality check of HMW DNA extracted and used for SMRTbell library preparation indicates A) large size fragments as shown by the TapeStation peak view and B) high molecular weight across all extractions.

### DNA Sequencing Library, Read filtering, and Genome Assembly

gDNA was sent to Genewiz (Azenta Life Sciences) for library preparation and sequencing using PacBio Sequel I long read technology. A single SMRTbell sequencing library (double-stranded DNA template with hairpins at each end) was constructed. Briefly, the DNA was fragmented by shearing, and DNA damage within the strand and at the ends of the fragments was repaired and the sample purified with AMPure PB beads. Hairpin adapters were then ligated to the DNA fragment ends and the sample again purified with AMPure PB beads. Sequencing primers were then annealed to form polymerase-primer complexes and the resulting library was sequenced using 4 SMRT cells on the PacBio Sequel I at GeneWiz (Table 1).

**Table 1.** Contig length and assembly statistics.

| Statistics | Filtered Assembly | Unfiltered Assembly |
| --- | --- | --- |
| No. contigs | 3,051 | 3,095 |
| No. contigs ($\geq 0$ bp) | 3,051 | 3,095 |
| No. contigs ($\geq 1000$ bp) | 3,051 | 3,095 |
| No. contigs ($\geq 5000$ bp) | 3,048 | 3,093 |
| No. contigs ($\geq 10000$ bp) | 3,030 | 3,070 |
| No. contigs ($\geq 25000$ bp) | 2,866 | 2,905 |
| No. contigs ($\geq 50000$ bp) | 2,429 | 2,432 |
| Largest contig (bp) | 3,369,715 | 2,514,820 |
| Total length (bp) | 677,753,397 | 680,005,989 |
| Total length ($\geq 0$ bp) | 677,753,397 | 680,005,989 |
| Total length ($\geq 1000$ bp) | 677,753,397 | 680,005,989 |
| Total length ($\geq 5000$ bp) | 677,742,019 | 679,999,105 |
| Total length ($\geq 10000$ bp) | 677,611,785 | 679,826,819 |
| Total length ($\geq 25000$ bp) | 674,350,533 | 676,554,116 |
| Total length ($\geq 50000$ bp) | 658,123,657 | 658,911,805 |
| N50 | 412,256 | 408,687 |
| N75 | 189,769 | 190,323 |
| L50 | 437 | 444 |
| L75 | 1,049 | 1,055 |
| GC (%) | 39.12% | 39.14% |

Prior to assembly, reads were mapped to the *Porites lutea* genome (Robbins et al. 2019) with MiniMap v2 (Li 2018) and SAMtools v1.9 (Li et al. 2009) to exclude non-coral reads. The filtered reads were assembled with Falcon v2.2.4 (Chin et al. 2016) at Genewiz. To assess genome assembly completeness, the *P. astreoides, Porites rus* (Celis et al. 2018), *P. lutea* (Robbins et al. 2019), and *Porites australiensis* (Shinzato et al. 2021) genomes were searched against 954 universal metazoan single-copy orthologs in the metazoa_odb10 gene set using BUSCO v5.2.2 in “genome” mode (Simão et al. 2015) (Table 2).

**Table 2.** Comparison of genome assembly and annotation metrics of Porites congeners.

| Feature | *Porites rus* | *Porites lutea* | *Porites australiensis* | *Porites astreoides* |
| --- | --- | --- | --- | --- |
| Study | [61] | [57] | [62] | This Study |
| Assembly size (Mbp) | 470 | 552 | 576 | 678 |
| No. protein coding genes | 39,453 | 31,126 | 30,301 | 64,636 |
| Mean no. of exons per gene | 5.3 | 6.6 | 7.6 | 4.9 |
| Mean exon length (bp) | 239 | 275 | 346 | 190 |
| Mean intron length (bp) | 1224 | 1146 | 1190 | 863 |
| Metazoan BUSCOs (%(gene number), total=954 genes) |  |  |  |  |
| Complete single-copy | 65.5 (625) | 91.9 (877) | 86.6 (826) | 77.5 (739) |
| Complete duplicated | 2.3 (22) | 1.8 (17) | 3.1 (30) | 13.4 (128) |
| Fragmented | 18.7 (178) | 2.8 (27) | 5.8 (55) | 4.1 (39) |
| Missing | 13.5 (129) | 3.5 (33) | 4.5 (43) | 5.0 (48) |

### Structural and Functional Annotation

Structural annotation of the *P. astreoides* genome was completed on the University of Rhode Island High Performance Computer “Andromeda”. As input for MAKER v3.01.03 (Cantarel et al. 2008) we used an existing *P. astreoides* transcriptome from samples collected in the Florida Keys, USA (Kenkel, Meyer, and Matz 2013) and existing congener *P. lutea* peptide sequences from a sample collected in Australia (Robbins et al. 2019). These files were used as input for an initial round of MAKER to predict gene models directly from this transcriptomic and protein data, respectively. The first round of MAKER included RepeatMasker (Chen 2004). The output from the initial MAKER round was used to train *ab initio* gene predictors SNAP (Korf 2004) and AUGUSTUS (Stanke et al. 2006) using BUSCO v5.2.2 (Simão et al. 2015). A second round of MAKER was performed using the GFF output of the first round containing the information on the locations of repetitive elements for masking, as well as the locations of ESTs and proteins and *ab initio* gene prediction from the SNAP and AUGUSTUS outputs. Another round of *ab initio* gene prediction was performed on the output from the second round of MAKER. A third and final round of MAKER was conducted including training information from the GFF files generated from the second round of MAKER and *ab initio* gene prediction as input. Genome structural annotations were compared against all four *Porites* congeners using AGAT v0.8.1 (Table 2).

The *ab initio* transcriptome generated from the final round of MAKER was compared against other *de novo P. astreiodes* transcriptome assemblies (Kenkel, Meyer, and Matz 2013; Mansour et al. 2016; Walker et al. 2019) by assessing statistics from BUSCO v5.2.2 in “transcriptome” mode referencing the metazoa_odb10 gene set (Simão et al. 2015) (Table 3). The protein set produced from the final round of MAKER was annotated by identifying homologous sequences using BLASTp (Basic Local Alignment Search tool, e-value cut-off = 1e-5) against a hierarchical approach of the following databases: 1) UniProt-SwissProt (The UniProt Consortium 2017) 2) UniProt-TrEMBL (The UniProt Consortium 2017) and 3) the NCBI “nr” database. From the BLASTp searches, BLAST2GO (Conesa et al. 2005) was used to characterize putative gene functionality in addition to InterProScan (Jones et al. 2014).

**Table 3.**
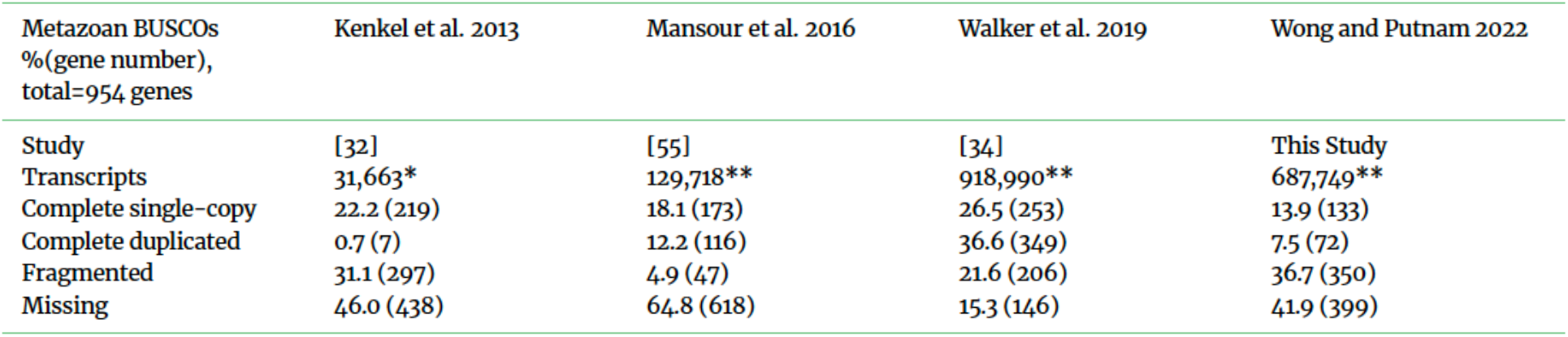
Comparison of BUSCO metrics of transcriptome completeness across *Porites astreoides* transcriptome assemblies relative to the BUSCO metazoa_odb10 gene set. Transcript quantification was conducted on isogroups using custom scripts for studies with a single asterisk (*) and on transcripts via Trinity for studies indicated with double asterisks (**).

### Extraction of Total RNA

Total RNA was extracted using the Duet DNA/RNA Miniprep Plus Kit (Zymo Research, Irvine, CA, USA, Cat #D7003) with modifications to the manufacturer’s sample preparation steps detailed here. Sterile clippers were used to remove ∼1cm diameter region from the coral tissue. The tissue clipping was placed in 500µL of DNA/RNA shield (Zymo Cat #R1100-50) with 0.5mm glass beads (Fisher Scientific Cat #15-340-152) and homogenized by vortexing for 1 minute. The supernatant (450µl) was transferred to a new tube and centrifuged at 9000 rcf for 3 minutes to remove any debris. The supernatant (300µl) was mixed with 30µl of Proteinase K (Zymo Cat #D3001-2-20) digestion buffer and 15µl of Proteinase K in a new tube using a brief vortex step. Equal volume of Zymo lysis buffer (Zymo Cat #D7001-1-200) was added (345µl) and extraction was completed as outlined in the manufacturer’s protocol. RNA concentration was quantified using the Qubit RNA Broad Range kit (Invitrogen Cat #Q10211) and integrity was assayed with an Agilent TapeStation 4200 system and the RNA Integrity Number (RIN) values ranged from 7.0-8.9. RNA libraries were prepared using the Zymo-Seq RiboFree Total RNA Library Kit (Zymo Cat #R3000) according to the manufacturer’s instructions starting with 125ng of RNA as input. Prepared libraries were sent to Zymo Research Corporation for sequencing using the Illumina HiSeq for paired end sequencing (2×100bp).

### RNASeq de novo Assembly and Genome Mapping

Four paired-end Illumina libraries generated 35,505,857 raw read pairs. The raw RNA-seq data was quality checked by using fastQC v0.11.8 (Andrews and Others 2010) and visualized with MultiQC v1.7 (Ewels et al. 2016). Illumina adapters were removed and sequences with a phred score below 30 were trimmed using fastp v0.19.7 (Chen et al. 2018). After trimming, 85-93% of the data was kept across the four libraries and the trimmed reads were used for *de novo* assembly and was completed using Trinity v2.9.1 (Grabherr et al. 2011) using default parameters except the fastq sequence type and CPU 20 parameters. The trimmed RNA-seq reads were also aligned to the *P. astreoides* genome using STAR v2.7.2b (Dobin et al. 2013).

## Results and Discussion

### Genome Assembly Statistics

The sequencing yielded just under 1,000,000 polymerase reads and a total polymerase read length of 20,188,576,450 (Table 1) from a single gDNA library. After quality control and assembly, we obtained a reference genome with a total size of ∼678Mb (Table 2). This *P. astreoides* genome had comparable assembly statistics to three other *Porites* species, *P. rus* (Celis et al. 2018) (assembly size = 470Mbp), *P. lutea* (Robbins et al. 2019) (assembly size = 552Mbp) and *P. australiensis* (Shinzato et al. 2021) (assembly size = 576Mbp) (Table 2).

### Genome Structural and Functional Annotation

The initial round of MAKER predicted 58,308 putative gene models, the second round 68,481 gene models, and 64,636 were predicted gene models in the third and final round. From our final structural annotation, *P. astreoides* had an average of 4.9 exons per gene, a mean exon length of 190bp, and a mean intron length of 863. This is comparable to other *Porites* congeners as shown in (Table 2). Of the 64,636 protein encoding genes, 86.6% (n= 55,957) in total were annotated with 47.1% (n= 30,444) receiving hits from the SwissProt database, 37.7% (n= 24,359) from the TrEMBL database, and 1.8% (n= 1154) from the NCBI “nr” database. 13.4% (n= 8,679) had no hits to any of the databases. Through BLAST2GO (Conesa et al. 2005), 30,284 genes were assigned putative protein function against the SwissProt database, and 24,089 were assigned with InterProScan (Jones et al. 2014). After removal of duplicate gene functions, 67% (n=43154) had assigned protein functions using BLAST2GO (Conesa et al. 2005) and InterProScan (Jones et al. 2014).

### Data Validation and Quality Control

Genome assembly completeness was assessed by searching 954 universal metazoan single-copy orthologs in the metazoa_odb10 gene set using BUSCO v5.2.2 (Simão et al. 2015). For the *P. astreoides* genome, 739 (77.5%) complete single-copy orthologs were identified, 128 (13.4%) orthologs were completed but duplicated, 39 (4.1%) orthologs were identified but fragmented, and 48 (5.0%) were missing. Compared to other *Porites* species, our *P. astreoides* assembly identified more complete single-copy orthologs than *P. rus* (Celis et al. 2018)(625 (65.5%) single-copy orthologs) but less than *P. lutea* (Robbins et al. 2019) (877 (91.9%) single-copy orthologs) and *P. australiensis* (Shinzato et al. 2021) (826 (86.6%) single-copy orthologs) (Table2). The BUSCO analysis includes 13.4 % duplication, which suggests the presence of haplotigs that were not fully removed in prior to or during the assembly process and should be addressed in future versions of the assembly.

To describe the mapping potential of the draft *P. astreoides* genome, four paired-end RNA-seq libraries were mapped to the *ab initio* reference genome using STAR v2.7.2b (Dobin et al. 2013). From the four RNA-seq libraries, mapping percentages ranged from 87.9% to 79.07%, with 17.9% to 17.05% of the reads mapping to multiple loci. This suggests that we have a suitable *ab initio* reference genome for RNA-seq data for Bermudian populations of *P. astreoides*.

Comparison of our *de novo* assembly statistics to those previously published (Table 3) indicates that all *de novo* assemblies to date suffer from transcript fragmentation or duplication in comparison to the BUSCO metazoan reference set. For example, even in the *de novo* transcriptome by Walker et al. (2019) with the highest complete and single copy BUSCO score and the lowest number of missing reference genes, the values are still only 26.5% complete and single copy and 15.3% remain missing.

While it is possible that there is insufficient gene model prediction in our draft assembly, the high number of gene models is likely to be a result of duplicated contigs from different haplotypes and a fragmented assembly, which is also seen currently in the *de novo* transcriptome. This issue of high predicted gene model number in *P. astreoides* should be improved in subsequent draft assemblies by purging haplotigs and the use of combined short read sequence data with our PacBio scaffolds. Here we have generated a valuable resource for the community that can now be improved through an iterative process through the coral research community investment.

### Reuse Potential

Due to the ecological increase in relative abundance of *P. astreoides* in the Atlantic (Green, Edmunds, and Carpenter 2008), it is crucial to understand the mechanisms leading to resilience under predicted climate change conditions. Given the potential to improve current transcriptomes (Kenkel, Meyer, and Matz 2013; Mansour et al. 2016; Walker et al. 2019) and the lack of genomic resources for this species, we provide the first draft reference genome for *P. astreoides*. Although we acknowledge this genome can and should be improved, the goal was to provide a community resource for the advancement of molecular biology and comparative genomics of reef-building corals. *P. astreoides* and the genus *Porites* in general is known to be a challenging coral species for molecular biology approaches. *Porites* species have high mucus (Johnston and Rohwer 2007) and lipid (Grottoli, Rodrigues, and Juarez 2004; Teece et al. 2011) content, thick tissues (Yost et al. 2013; Putnam et al. 2017), and high endosymbiont densities (Thornhill et al. 2011), which can create complications for DNA and RNA extractions and retain molecules that act as inhibitors in PCR, and thus often result in poor quality library preparation and *de novo* transcriptome assemblies. The *ab initio* reference genome, transcriptome, and updated annotations generated by this study therefore have the potential to serve as a useful resource to the coral field and marine invertebrate community more broadly.

### Availability of supporting data and source code

The genome assembly and all raw sequencing reads including the PacBio HiFi reads and Illumina RNA-seq, annotations, alignments, and other results are available via Open Science Framework Repository DOI 10.17605/OSF.IO/ED8XU (https://osf.io/ed8xu/).

## List of abbreviations

DNA: Deoxyribonucleic acid
RNA: Ribonucleic acid
gDNA: Genomic DNA
PCR: Polymerase Chain Reaction
BLAST: Basic Local Alignment Search Tool
bp: base pair
Gbp: gigabase pair
Mbp: megabase pair
kbp: kilobase pair
HMW: high molecular weight
BUSCO: Benchmarking Universal Single-Copy Orthologs
NCBI: National Center for Biotechnology Information

## Ethical Approval

Corals were collected under permit 17060807 from the Government of Bermuda Department of Environment and Natural Resources and exported with CITES permit number 19BM0011.

## Author’s Contributions

Conceptualization: HMP, Methodology HMP, KHW; Investigation: KHW, HMP; Formal analysis: KHW; Resources: HMP; Data Curation: KHW, HMP; Writing Original Draft: KHW, HMP; Writing Review & Editing: KHW, HMP; Visualization: KHW; Supervision: HMP; Funding acquisition: HMP

## Acknowledgements

We would like to acknowledge the Bermuda Institute of Ocean Sciences facilities, Heising-Simons Foundation International, LTD, Natural Sciences and Engineering Research Council of Canada Post Graduate Scholarship Doctoral (NSERC PGSD3 Award #545967-2020), University of Rhode Island High Performance Computing, and the Canadian Associates of the Bermuda Institute of Ocean Sciences for their logistical and financial support of this work. Additionally, we would like to thank Dr. Samantha de Putron, Dr. Gretchen Goodbody-Gringley and Alexander Chequer for field assistance, Margaret Schedl for laboratory assistance and Kevin Bryan at the University of Rhode Island High Performance Computing for his assistance with computational needs.

